# DNA methylation estimation using methylation-sensitive restriction enzyme bisulfite sequencing (MREBS)

**DOI:** 10.1101/217208

**Authors:** Giancarlo Bonora, Liudmilla Rubbi, Marco Morselli, Constantinos Chronis, Kathrin Plath, Matteo Pellegrini

## Abstract

Whole-genome bisulfite sequencing (WGBS) and reduced representation bisulfite sequencing (RRBS) are widely used for measuring DNA methylation levels on a genome-wide scale(1). Both methods have limitations: WGBS is expensive and prohibitive for most large-scale projects; RRBS only interrogates 6-12% of the CpGs in the human genome(16,19). Here, we introduce methylation-sensitive restriction enzyme bisulfite sequencing (MREBS) which has the reduced sequencing requirements of RRBS, but significantly expands the coverage of CpG sites in the genome. We built a multiple regression model that combines the two features of MREBS: the bisulfite conversion ratios of single cytosines (as in WGBS and RRBS) as well as the number of reads that cover each locus (as in MRE-seq(12)). This combined approach allowed us to estimate differential methylation across 60% of the genome using read count data alone, and where counts were sufficiently high in both samples (about 1.5% of the genome), our estimates were significantly improved by the single CpG conversion information. We show that differential DNA methylation values based on MREBS data correlate well with those based on WGBS and RRBS. This newly developed technique combines the sequencing cost of RRBS and DNA methylation estimates on a portion of the genome similar to WGBS, making it ideal for large-scale projects of mammalian genomes.

## INTRODUCTION

DNA methylation plays an important role in gene regulation and the maintenance of cell identity, although much remains to be uncovered regarding the specific mechanisms underlying the targeting and reading of methylcytosines(2-4). High-throughput DNA sequencing technologies have enabled the measurement of cytosine methylation on a genome-wide scale, leading to the application of these approaches in myriad studies(5-12). Many technologies have been developed over the past decade to measure DNA methylation(13). Some of these provide qualitative information about regions enriched for DNA methylation, such as approaches that select DNA fragments using proteins that selectively bind methylated cytosines (e.g. meDIP)(13,14), or methods to digest DNA with methylation-sensitive restriction enzymes (e.g. MRE)(15). Other approaches are able to probe the methylation state of single cytosines, by chemically converting unmethylated cytosines to uracils using sodium bisulfite(16). The fraction of unconverted cytosines provides an estimate of the DNA methylation level at a particular locus. This read out can be determined from microarrays (e.g. Illumina 450K)(17) or next-generation sequencing(13,18-20). In addition, some single molecule sequencing technologies are able to directly detect methylcytosines by monitoring the incorporation rates of nucleotides (e.g. Pacific Biosciences)(21).

Among these approaches, two widely used next-generation sequencing methods for assessing DNA methylation levels at single cytosines on a genome-wide scale are whole-genome bisulfite sequencing (WGBS, also known as BS-seq, methyl-seq, or methylC-seq)(18,19,22) and reduced representation bisulfite sequencing (RRBS)(20,23). As implied by their names, both protocols are based on bisulfite treatment of DNA. However, to obtain high-confidence methylation estimates, one requires a minimum level of read coverage per site, typically at least 5–10X. WGBS is far more comprehensive and can in theory assess the methylation status of nearly every single cytosine in the genome, but requires very deep sequencing to arrive at modest levels of coverage, and, hence, can be very costly, especially if one is working with large genomes, such as those of mammals.

RRBS interrogates a smaller portion of the genome, significantly reducing the amount of sequencing required to obtain high-confidence methylation estimates at this subset of sites. The RRBS protocol introduces a step where genomic DNA is first digested, typically with the methylation-insensitive restriction endonuclease MspI, which cuts at the recognition sequence CˇCGG. Digested fragments are then size selected, typically in the range of 50 to 300 nucleotides. This fraction enriches for CpG-rich regions, including many regions involved in transcriptional regulation such as promoters and enhancers, but typically only covers 6–12% of CpGs genome-wide(23,24).

To address the respective limitations of WGBS and RRBS, we developed a new method, methylation-sensitive restriction enzyme bisulfite sequencing (MREBS), which adds a bisulfite step to an existing protocol: MRE-seq(15). Typically, MRE-seq utilizes three methylation-sensitive restriction endonucleases in parallel to digest DNA (HpaII (CˇCGG), HinP1I (GˇCGC), and AciI (CˇCGC)). Similarly to RRBS, a size selection step enriches for fragments between 50 bp and 300 bp. DNA methylation levels are inferred by the inverse relationship between MRE-seq read coverage and CpG methylation at the restriction enzyme target sites. Although the three aforementioned restriction enzymes only cut DNA at unmethylated CpG dinucleotides, the DNA methylation state of other CpGs within the resulting fragments could still be either methylated or unmethylated. We reasoned that the addition of a bisulfite conversion step to the MRE-seq protocol would directly measure the methylation state of cytosines within MRE fragments. Typically 70 – 80 % of CpG dinucleotides in the genome are methylated(2), and the rationale behind this approach is that we direct our sequencing resources to the regions of the genome that are more likely to be unmethylated by using MRE to digest the DNA. Then, rather than simply relying on the inverse relationship between MRE-seq read coverage and DNA methylation levels around cut sites, we additionally directly measure the DNA methylation levels of their flanking regions. In principle, the advantage afforded by MREBS over WGBS and RRBS, is that we focus our sequencing effort on hypomethylated loci.

Since MREBS reads are expected to show an overall bias for lowly methylated regions due to the propensity of the restriction enzymes to digest demethylated regions, their methylation levels do not provide an unbiased measurement of absolute DNA methylation levels. However, the data can be readily used to determine differential methylation between two samples, which is often of greater interest. With this in mind, we developed a computational model that determines differential DNA methylation in two ways. First, based on read coverage alone, which is expected to anti-correlate with DNA methylation levels, we determined differential methylation within a region around each CpG dimer by looking at the difference in read counts between samples, as is done with traditional MRE-seq. Second, in those regions with sufficient read coverage for reliable estimates, we determined differential DNA methylation at single CpG resolution based on a model of bisulfite conversion ratios.

To test our approach, we first compared MREBS conversion-based methylation estimates and coverage to those based on WGBS and RRBS data, using two cell types that represent very different developmental stages, namely mouse embryonic stem cells (ESCs) and an early somatic cell reprogramming intermediate obtained by inducing the expression of Oct4, Sox2, Klf4, and cMyc in mouse embryonic fibroblasts (MEFs) for 48 hours, where we had observed substantial differential methylation by WGBS and RRBS. We found that MREBS bisulfite conversion-based DNA methylation estimates correlated well with WGBS and RRBS-based values. The number of CpG dimers with sufficient read coverage to obtain MREBS conversion-based methylation estimates was comparable to that of RRBS. Importantly, in contrast to RRBS, we found that nearly 60% of all CpGs in the mouse genome had sufficient reads within the surrounding region in at least one of the two cell types to determine differential DNA methylation estimates based on MREBS read counts alone. Within lower coverage regions, we compute the counts in 1kb windows around CpGs to obtain approximate differential methylation as with traditional MRE-seq. In high coverage regions (~3% of CpGs), we use a multiple regression model that considers both cytosine methylation estimates from converted reads and read count data to predict differential DNA methylation values. The differential methylation estimates generated by this model compared favorably to measurements from RRBS data.

We found that MREBS provides a level of sequence coverage with nucleotide resolution similar to that obtained with RRBS. Additionally, with MREBS one can estimate DNA methylation levels for broader swathes of the genome based on differential MRE read counts around CpGs, thereby providing a level of coverage that begins to approach that obtained by WGBS, but at a fraction of the cost.

## MATERIAL AND METHODS

### Methylation-sensitive restriction enzyme bisulfite sequencing (MREBS)

Three enzymatic digestions were performed on 1 μg of purified genomic DNA using 10 U of each one of the MRE restriction enzymes (HpaII, Hin6 and AciI - Fermentas) in a 50 μl final volume with TANGO buffer. 2.5 μl of RNase cocktail mix (Ambion) were added and the reaction was incubated overnight at 37°C. After the digestion, the three reactions were pooled and the DNA was purified using AMPure XP beads (Beckman Coulter). Subsequent reactions of DNA End Repair, A-tailing and Adapter Ligation were performed using Illumina TruSeq reagents, following manufacturer’s instructions and the DNA was size selected between 200 and 500 bp using AMPure XP beads. Size selected DNA was then treated with bisulfite using the EpiTect kit (QIAGEN) according to the protocol suggested from the manufacturer, except that the conversion step was performed twice, for a total time of 10 h. For each bisulfite-converted sample, two parallel PCR reactions were set up in a final volume of 50 μl using MyTaq HS Mix (Bioline) and 2.5 μl of Illumina TruSeq PCR Cocktail Primers. The amplification cycles were as follows: 98°C – 2 min; 12 cycles of: 98°C – 15 sec, 60°C – 30 sec, 72°C – 30 sec; 72°C – 5 min. The final PCR products were purified using AMPure XP beads and the final concentration of the libraries was measured using Qubit DNA BR Assay (Life Technologies). Single-end sequencing for 100bp reads was performed on an Illumina Hiseq 2000.

### Whole-genome bisulfite sequencing (WGBS)

Genomic DNA from induced MEFs (48h OSKM) and ESCs was isolated using the Blood and tissue DNeasy kit (Qiagen). Isolated DNA was treated with RNAseA for 30 min at 37oC and cleaned up using AMPure XP beads. 5 μg of treated DNA was fragmented to 100–500 bp using a Bioruptor Sonicator. 5 minutes in pulses of 30 sec on, 1 minute off. DNA fragments were visualized on 1% agarose gel, gel extracted and purified using a QIAGEN gel extraction minelute kit. End-repair reactions (50 μl) contained 1x T4 DNA ligase buffer (NEB), ATP, 0.4 mM dNTPs, 15 units T4 DNA polymerase, 5 units Klenow DNA polymerase, 50 units T4 polynucleotide kinase (all NEB) and were incubated for 30 min at 20°C. DNA clean-up was performed using a 2x volume of AMPure XP beads and eluted in 32 μl of dH2O. Adenylation was performed for 30 minutes at 37°C in 50 μl volumes that contained 5 μl 1x Klenow buffer, 0.2 mM dATP and 15 units Klenow exo− (NEB). Adenylated DNA fragments and methylated adapters (Illumina) were ligated for 15 min at 20oC in a 50 μl reaction containing 5,000 units quick ligase (NEB) and 5 μl of adapters. Adaptor-ligated DNA of 200-600 bp, was size-selected on a 2% agarose gel. Bisulfite conversion was performed with an EpiTect Bisulfite Kit (QIAGEN) following the manufactures conditions. Bisulfite converted DNA was amplified for 15 cycles with PfuTurboCx Hotstart DNA polymerase (Agilent technologies). The final library DNA was quantified using a Qubit fluorometer and a Quant-iT dsDNA HS Kit (Invitrogen). Single-end sequencing for 100bp reads was performed on an Illumina Hiseq 2000.

### Reduced Representation Bisulfite sequencing (RRBS)

5 μl of genomic DNA was digested with 50 units of MspI (NEB) in a 100 μl reaction for 6 hours at 37oC. Digested DNA was run on a 3% low-melt agarose gel (Lonza) and fragments of 25 to 300 bp were extracted and purified using a MinElute gel extraction kit (QIAGEN) according to the manufacturers instructions. DNA end-repair and adenylation was as described above with the exception of using a dNTP mix consistent of dATP, dGTP and 5medCTP. Ligation to methylated adapters and subsequent library construction was performed similarly to the WGBS protocol. Single-end sequencing for 100bp reads was performed on an Illumina Hiseq 2000.

### Bisulfite sequencing data processing

DNA methylation calling was performed using BS-Seeker2(27) using Bowtie 0.12.9(30) for read alignment. WGBS and MREBS reads were mapped to the mm9 reference genome while RRBS reads were mapped to a reduced reference that was in silico digested using the MspI recognition sequence and limited to fragments of 20–500bp in length. The 100bp reads were trimmed of adapter sequences and allowed 5 mismatches during mapping. MREBS reads were first filtered so that only the expected 5’ trimers (CGG and CGC; Supplementary Table 1) were retained. For conversion based DNA methylation level calling, only CpG dimers covered by at least 5 reads on both were used in an effort to obtain reliable methylation levels.

### ChIP-seq library preparation and chromatin states analysis

The protocol and the model is described in detail in a separate manuscript(26), but briefly 18 chromatin states in the MEFs, EARLY intermediates, LATE intermediates, and ESCs were identified at a resolution of 200 bp using chromHMM as described by Ernst and Kellis(31) using ChIP-seq data sets for nine histone modification, one histone variant (H3.3), and an input, as listed in Figure 2.

**Figure 2.**
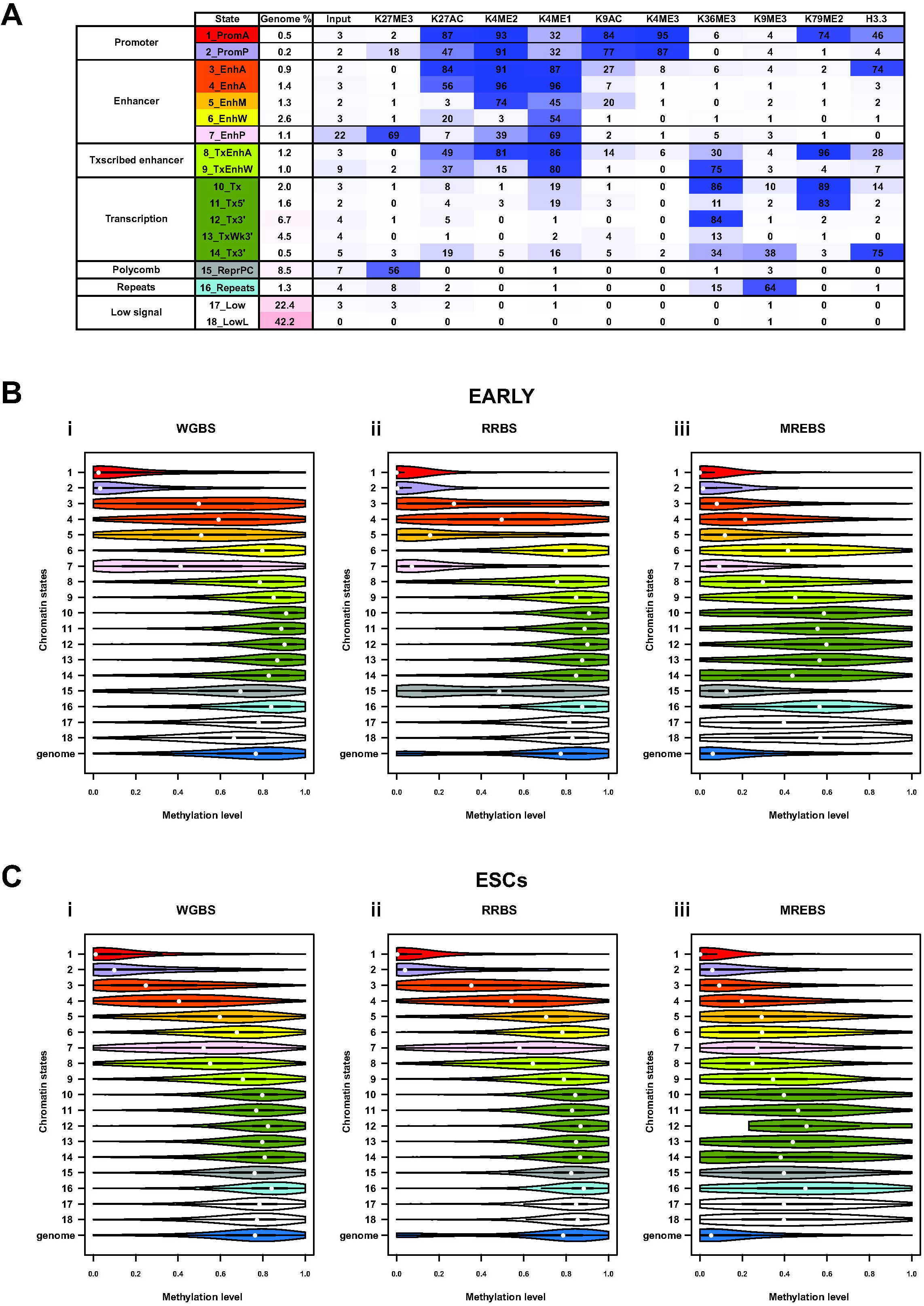
DNA methylation estimates based on WGBS, RRBS and MREBS data in different chromatin states. A. Heat map and functional annotation for an 18-level chromatin state model at 200 bp resolution built using peak calls made using ChIP-seq data sets for 10 histone modifications, as well as an input library, for each of the three cell types described in Figure 1A: i) MEFs, ii) EARLY intermediates, and iii) ESCs, as well as a late reprogramming intermediate (LATE), partially induced pluripotent cells, or pre-iPSCs, not otherwise used in the study. Candidate functional annotations were assigned to each of the 18 chromatins states based on the prevalence and combination of histone mark peaks, which could in turn be classified into the seven categories indicated in the left-hand column. The probability of a window in each state to be contain a peak for a given histone modification is given as a percentage in each cell, and visually indicated by the intensity of color in the heat map. The proportion of the concatenated genome (MEFs + EARLY + LATE + ESCs) found in each of the 18 chromatin states is given in the third column. B. Violin plots of the distributions of the DNA methylation estimates in each of the 18 chromatin states (described in A), as well as genome-wide (blue), for EARLY intermediates using i) whole-genome bisulfite sequencing (WGBS), reduced representation bisulfite sequencing (RRBS), and iii) methylation-sensitive restriction enzyme bisulfite sequencing (MREBS). The mean DNA methylation estimates within 200 bp windows corresponding to those used for the chromatin state model were used, only considering those windows containing at least one CpG with 5X coverage, in an effort to ensure high-confidence estimates. White circles represent median values. C. As in (B), but for ESCs.

### Differential DNA methylation modeling

Linear regression was used to model differential CpG dimer methylation estimates based on WGBS (the response vectors) using differential methylation estimates based on RRBS and MREBS, as well as differential read counts within 1kb windows based on MREBS data around corresponding CpG dimers with R's lm() function(29). The coefficients in in Supplementary Table 5 are outputs from R's summary.lm() function(29).

### Data submission

All high-throughput sequencing data will be made available on GEO at the time of publication.

## RESULTS

### Study design and data sets

We chose to test our approach on two cell lines where we expected to see significant differential methylation as they represented distinct developmental stages and differentially methylated regions (DMRs) were observed using WGBS and RRBS. The two cell types were: 1) MEFs that were induced to ectopically express the Yamanaka reprogramming factors OCT4 (O), SOX2 (S), KLF4 (K), and MYC (M; also known as ‘cMYC’)(25) for 48 hours, representing an early somatic cell reprogramming intermediate (EARLY), and 2) mouse embryonic stem cells (ESCs) representing the pluripotent stem cell state reached upon successful reprogramming (Figure 1A). Both states have been recently described in detail (26). WGBS libraries for the two cell types were generated and reads were mapped to the mm9 genome using BS-Seeker2(27). The WGBS data sets for the two cell types showed comparable sequencing depth and CpG coverage (Figure 1B, Supplementary Table 2). RRBS and MREBS libraries for the same cell lines were also generated, with the MREBS libraries produced in duplicate to test reproducibility. RRBS reads were mapped to an in silico MspI-digested reduced reference mm9, and MREBS reads were mapped to the whole genome after being filtered for the expected 5’ cut sites (Figure 1C/D, Supplementary Table 1). Although the sequencing depth for RRBS and MREBS was comparable, twice as many CpGs were covered by at least one read with MREBS (Figure 1C/D, Supplementary Table 2).

**Figure 1.**
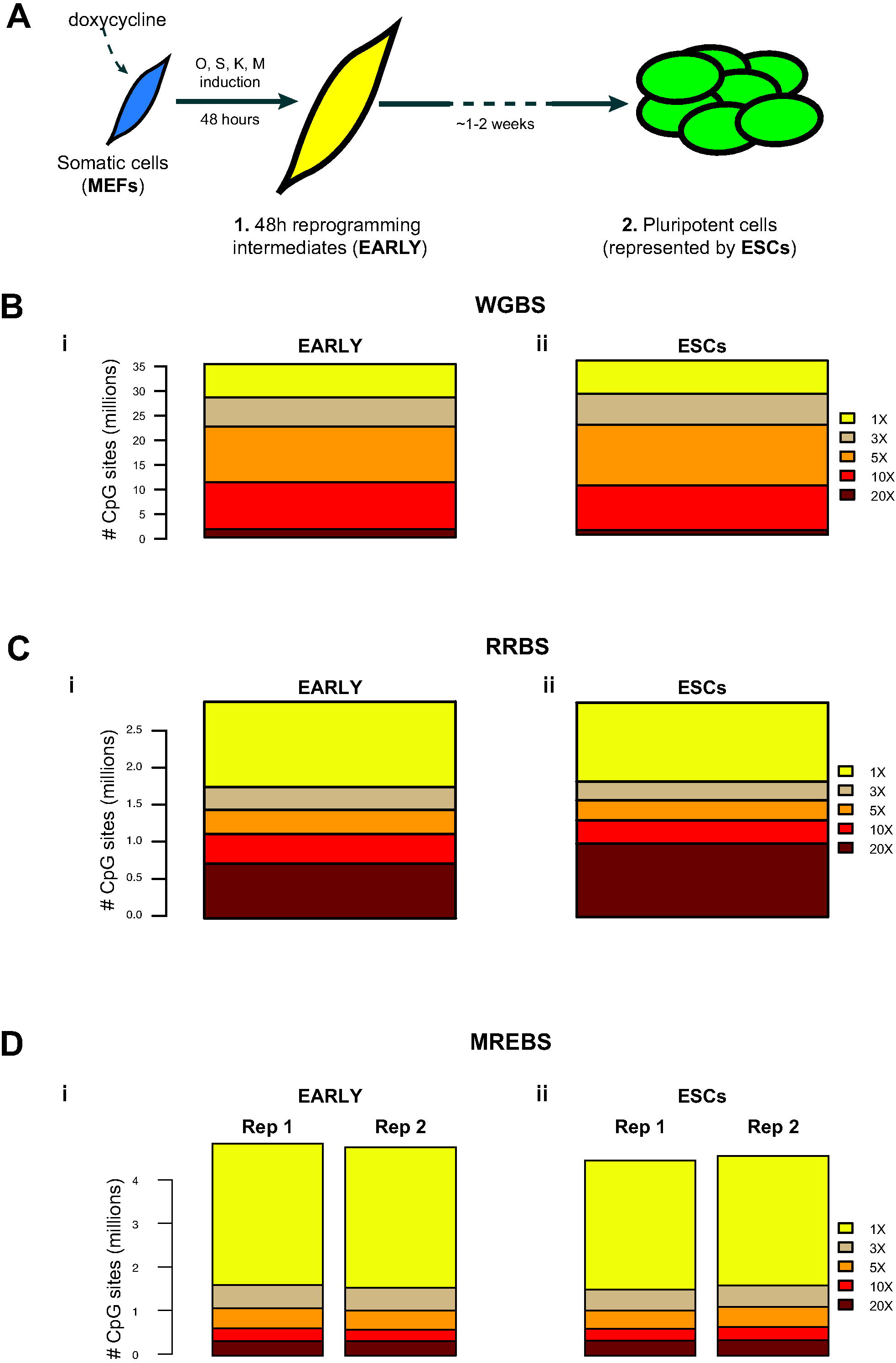
WGBS, RRBS, and MREBS for samples representing two stages of somatic cell reprogramming. A. Schematic representation of the two cell types used in the study. Mouse embryonic fibroblasts (MEFs; blue), modified to harbor a ‘stem cell cassette’ allowing for the simultaneous induction of the four pluripotency factors (OCT4 (O), SOX2 (S), KLF4 (K), and MYC (M)) by the addition of doxycycline, were induced for 48 hours. These EARLY somatic cell reprogramming intermediates (yellow) were the first of the two cell types sampled, with embryonic stem cells (ESCs; green), representing the fully reprogrammed state, being the second. B. Bar plots showing the number of CpGs obtained at five different coverage levels (1–20 X) in each of the two whole-genome bisulfite sequencing (WGBS) samples: i) EARLY intermediates (429 M mapped reads) and ii) ESCs (391 M mapped reads). C. As in (B), but for reduced representation bisulfite sequencing (RRBS) samples (12.8 M mapped reads for i and 18.1 M mapped reads for ii). D. As in (B), but for duplicate samples produced using methylation-sensitive restriction enzyme bisulfite sequencing (MREBS) (11.9 and 12.4 M mapped reads for i; 11.8 and12.2 M mapped reads for ii).

### MREBS read counts provide high coverage of the genome

To determine DNA methylation estimates using bisulfite conversion rates, one typically requires at least 5X read coverage. As expected, the proportion of the 21.3 million CpGs in the mouse genome covered with a minimum of 5X coverage was substantially higher for the WGBS samples (~80%), than for either the RRBS (6%) or the MREBS samples (4–5%) (Supplementary Table 3). And, as was the case for the individual samples, the pairwise 5X coverage was substantially higher for the WGBS samples (75.5% of all CpG dimers between the two samples) than for either the RRBS (5.6%) or the MREBS samples (~3%) (Supplementary Table 4). However, apart from using bisulfite conversion ratios to determine DNA methylation, we reasoned that for the MREBS samples we might be able to model DNA methylation based on differential read coverage alone as with traditional MRE since MREBS utilizes methylation sensitive digestion, read counts around each CpGs should anti-correlate with their methylation levels (Tables 1 and 2). In other words, MREBS read counts within windows around CpGs could be used to determine methylation, thereby providing broader coverage than one would obtain by relying only on high confidence DNA methylation calls at each CpG based on bisulfite conversion ratios. 42–48% of CpG dimers had two or more reads falling within a surrounding 1kb window (Supplementary Table 3), with nearly 60% of CpG dimers had at least two MREBS reads falling within the surrounding 1kb window in at least one of the two cell types (Supplementary Table 4). This suggested that MREBS could be utilized for determining differentially methylated regions (DMRs) between a pair of samples using both read counts and bisulfite conversion ratios.

**Table 1.** CpG dimer-level correlations between bisulfite sequencing libraries. Pearson correlation values between WGBS, RRBS, and MREBS CpG dimer-level conversion-based methylation estimates, as well as binned read counts within 1kb windows around CpG dimers, for those with at least two MREBS reads. Red intensity signifies the strength of a positive correlation, while blue intensity signifies the strength of the anti-correlation.

|  |  |  | CpG dimer-level conversion-based methylation estimates * |  |  |  |  |  |  |  | MREBS read counts within 1kb windows around CpG dimers ** |  |  |  |
| --- | --- | --- | --- | --- | --- | --- | --- | --- | --- | --- | --- | --- | --- | --- |
|  |  |  | WGBS |  | RRBS |  | MREBS |  |  |  |  |  |  |  |
|  |  |  | EARLY | ESC | EARLY | ESC | EARLY Rep1 | EARLY Rep2 | ESC Rep1 | ESC Rep2 | EARLY Rep1 | EARLY Rep2 | ESC Rep1 | ESC Rep2 |
| CpG dimer-level conversion-based methylation estimates * | WGBS | EARLY | 1.00 | 0.56 | 0.90 | 0.78 | 0.78 | 0.77 | 0.55 | 0.55 | -0.14 | -0.14 | -0.18 | -0.17 |
|  |  | ESC | 0.56 | 1.00 | 0.77 | 0.90 | 0.53 | 0.53 | 0.70 | 0.71 | -0.15 | -0.15 | -0.24 | -0.22 |
|  | RRBS | EARLY | 0.90 | 0.77 | 1.00 | 0.88 | 0.82 | 0.82 | 0.66 | 0.66 | -0.21 | -0.21 | -0.27 | -0.27 |
|  |  | ESC | 0.78 | 0.90 | 0.88 | 1.00 | 0.64 | 0.64 | 0.80 | 0.80 | -0.21 | -0.21 | -0.30 | -0.29 |
|  | MREBS | EARLY Rep1 | 0.78 | 0.53 | 0.82 | 0.64 | 1.00 | 0.88 | 0.68 | 0.67 | -0.07 | -0.07 | -0.09 | -0.09 |
|  |  | EARLY Rep2 | 0.77 | 0.53 | 0.82 | 0.64 | 0.88 | 1.00 | 0.69 | 0.68 | -0.07 | -0.07 | -0.10 | -0.09 |
|  |  | ESC Rep1 | 0.55 | 0.70 | 0.66 | 0.80 | 0.68 | 0.69 | 1.00 | 0.83 | -0.07 | -0.07 | -0.10 | -0.10 |
|  |  | ESC Rep2 | 0.55 | 0.71 | 0.66 | 0.80 | 0.67 | 0.68 | 0.83 | 1.00 | -0.07 | -0.07 | -0.10 | -0.09 |
| MREBS read counts within 1kb windows around CpG dimers ** | EARLY Rep1 |  | -0.14 | -0.15 | -0.21 | -0.21 | -0.07 | -0.07 | -0.07 | -0.07 | 1.00 | 1.00 | 0.96 | 0.98 |
|  | EARLY Rep2 |  | -0.14 | -0.15 | -0.21 | -0.21 | -0.07 | -0.07 | -0.07 | -0.07 | 1.00 | 1.00 | 0.97 | 0.98 |
|  | ESC Rep1 |  | -0.18 | -0.24 | -0.27 | -0.30 | -0.09 | -0.10 | -0.10 | -0.10 | 0.96 | 0.97 | 1.00 | 0.99 |
|  | ESC Rep2 |  | -0.17 | -0.22 | -0.27 | -0.29 | -0.09 | -0.09 | -0.10 | -0.09 | 0.98 | 0.98 | 0.99 | 1.00 |

\* For CpG dimers with 5X coverage in both cell lines
\*\* For windows with at least two MREBS reads in at least one cell line

**Table 2.** CpG dimer-level correlations between differential values for all bisulfite sequencing library pairs. Pearson correlation values between WGBS, RRBS, and MREBS differential CpG dimer-level methylation estimates (EARLY - ESC), as well as differential read counts between all CpG dimers with at least two MREBS reads within a surrounding 1kb window. Red intensity signifies the strength of a positive correlation, while blue intensity signifies the strength of the anti-correlation.

|  |  |  | CpG dimer-level differential methylation using conversion-based estimates * |  |  |  | Differential MREBS read counts within 1kb windows around CpG dimers ** |  |
| --- | --- | --- | --- | --- | --- | --- | --- | --- |
|  |  |  | WGBS | RRBS | MREBS |  |  |  |
|  |  |  |  |  | Rep1 | Rep2 | Rep1 | Rep2 |
| CpG dimer-level differential methylation using conversion-based estimates * | WGBS |  | 1.00 | 0.63 | 0.55 | 0.56 | -0.34 | -0.34 |
|  | RRBS |  | 0.63 | 1.00 | 0.64 | 0.64 | -0.51 | -0.52 |
|  | MREBS | Rep1 | 0.55 | 0.64 | 1.00 | 0.64 | -0.35 | -0.35 |
|  |  | Rep2 | 0.56 | 0.64 | 0.64 | 1.00 | -0.34 | -0.34 |
| Differential MREBS read counts within 1kb windows around CpG dimers ** |  | Rep1 | -0.34 | -0.51 | -0.35 | -0.34 | 1.00 | 0.87 |
|  |  | Rep2 | -0.34 | -0.52 | -0.35 | -0.34 | 0.87 | 1.00 |

\* For CpG dimers with 5X coverage in both cell lines
\*\* For windows with at least two MREBS reads in at least one cell line

### MREBS conversion ratios correlate and MREBS read counts anti-correlate with WGBS and RRBS DNA methylation estimates, respectively

To investigate the relationship between WGBS, RRBS and MREBS-based DNA methylation estimates, we computed global correlations between them (Table 1). MREBS bisulfite conversion-based methylation estimates correlated more closely with the WGBS and RRBS-based estimates in the EARLY reprogramming intermediate, than they did with the ESC counterparts. Moreover, MREBS conversion-based methylation estimates for ESCs correlated more closely with the ESC RRBS and WGBS data than they did with the methylation estimates of EARLY reprogramming intermediate (Table 1). As expected, MREBS read counts anti-correlated with the DNA methylation levels based on WGBS-, RRBS-, and MREBS data, though not in a particularly cell type-specific manner (Table 1). Correlations based on differential data were substantially stronger than those based on absolute levels. MREBS differential data correlated with the differential WGBS and RRBS differential DNA methylation in the expected directions: differential read counts between the EARLY reprogramming intermediate and ESCs (EARLY - ESC) based on MREBS anti-correlated strongly with the differential DNA methylation values based on WGBS and RRBS, while MREBS differential bisulfite conversion ratio estimates correlated positively with those of WGBS and RRBS (Table 2). This suggested that MREBS data might be best utilized to estimate differential DNA methylation. Additionally, based on these observations, we hypothesized that by combining the MREBS differential conversion ratio estimates and MREBS differential read counts, we could make use of both domains of MREBS data to better estimate differential DNA methylation.

### Distributions of MREBS bisulfite conversion-based DNA methylation estimates across different chromatin states are similar to those of WGBS and RRBS-based estimates

To ensure that DNA methylation estimates based on MREBS data corresponded to those based on WGBS and RRBS in all genomic contexts, we compared DNA methylation estimates across different chromatin states. To determine chromatin states, we took advantage of a hidden Markov model of chromatin states generated by using chromHMM(28). The model is described in detail in Chronis et al.(26). Briefly, the genomes of the two cell types were tiled into 200 bp windows and assigned to one of 18 chromatin states based on ChIP-seq signals for nine histone modifications and one histone variant (histone H3.3), including a native input library. Functional annotations were determined for each of the 18 states based on the prevalence and combination of the histone mark peaks and the enrichment of genomic features (Figure 2A).

Mean DNA methylation levels were estimated for all those 200 bp windows containing a minimum of one CpG with 5X read coverage. Distributions of DNA methylation levels genome-wide and within the 200 bp windows belonging to each chromatin state were then plotted for each cell type (EARLY intermediates (Figure 2B) and ESCs (Figure 2C)) and for each bisulfite sequencing method (WGBS (i), RRBS (ii), and MREBS (iii)). Different chromatin states showed characteristic DNA methylation distributions that were similar in both cell types (Figure 2B/C). For instance, the promoter-associated chromatin states (1 and 2) were comparatively hypomethylated, while several of the enhancer-related chromatin states (3, 4, 5, and 7) showed wide spread DNA methylation levels and an intermediate mean DNA methylation. Most of the other chromatin states were largely hypermethylated (Figure 2B/C).

Apart from differences across chromatin states, there were also some differences between the cell types, as well as differences between approaches. For instance, the distributions of the MREBS-based DNA methylation estimates are systematically lower in most chromatin states as might be expected due to the use of methylation-sensitive endonucleases (Figure 2Biii/Ciii). Most notably the genome-wide DNA methylation levels based on MREBS estimates are low, (dark blue violin plots) close to that of the more demethylated chromatin states. Indeed, the MREBS samples are particularly enriched for two states of regulatory importance having the lowest DNA methylation levels, namely promoter and specific enhancer states (chromatin states 1–5, 7, and 15, Figure 2B/C). Although MREBS conversion-based DNA methylation estimates are systematically lower than those obtained by WGBS and RRBS data, their distributions within different chromatin states are very similar.

Reassured that MREBS DNA methylation estimates mirrored patterns see by WGBS and RRBS across all chromatins states, we hypothesize that MREBS could be used to determine methylation levels in and between samples, if scaled appropriately, or incorporated into a model to predict differential DNA methylation.

### Differential CpG-level methylation can be modeled using MREBS data

We hypothesized that WGBS-based differential DNA methylation values could be modeled using the MREBS data. To investigate this, we built four different linear regression models, as follows:

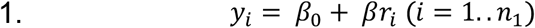

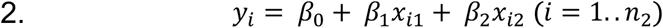

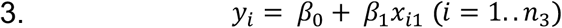

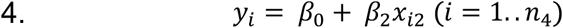

Here, y represents the WGBS conversion-based differential DNA methylation (EARLY - ESC), r the RRBS conversion-based differential DNA methylation, and x1 the MREBS conversion-based differential DNA methylation, in each case for those CpG dimers with 5X coverage in both cell types. x2 represents the differential MREBS read count within a 1kb window around each CpG dimer for windows with at least 2 reads in at least one of the two cell types. n1 = 666,214; n2 = 318,400 for MREBS replicate 1 and n2 = 322,431 for MREBS replicate 2; n3 = 319,304 for MREBS replicate 1 and n3 = 323,431 for MREBS replicate 2; n4 = 9,485,471 for MREBS replicate 1 and n4 = 9,670,440 for MREBS replicate 2.

In other words, in each case WGBS-based differential methylation serves as the response variable. In model 1, RRBS-based differential methylation is used as the explanatory variable. Model 2 is a multiple linear regression model that uses both MREBS conversion-based differential methylation and MREBS-based differential reads counts to predict WGBS-based differential methylation estimates, while model 3 and model 4 use each of these predictors independently. The lm() function from the R statistical software environment was used to implement these models(29). Coefficients for each model are provided in Supplementary Table 5. Model 1 was used for comparison purposes and shows that WGBS-based differential DNA methylation values (EARLY - ESC) can be modeled using RRBS data for ~3% of the CpG dimers (666,214) in the mouse genome with sufficient coverage (5X) in both the WGBS and RRBS samples. The model had an R2 = 0.39, implying a correlation between the WGBS estimated differential DNA methylation values and the model-fitted ones of r = 0.63 (Table 3). The root-mean-square error (RMSE) between the observed and fitted values was 20.4% and the mean absolute error (MAE) was 15.2%. The metrics ‘methyl15’ and ‘methyl25’ give the percentage of CpG dimers where the difference between the WGBS differential DNA methylation estimate and that of the model was at most 15% and 25%, respectively. Based on the methyl25 metric, the RRBS-based model-fitted DNA methylation values show 80% concordance with the WGBS-based estimates (Table 3).

**Table 3.**
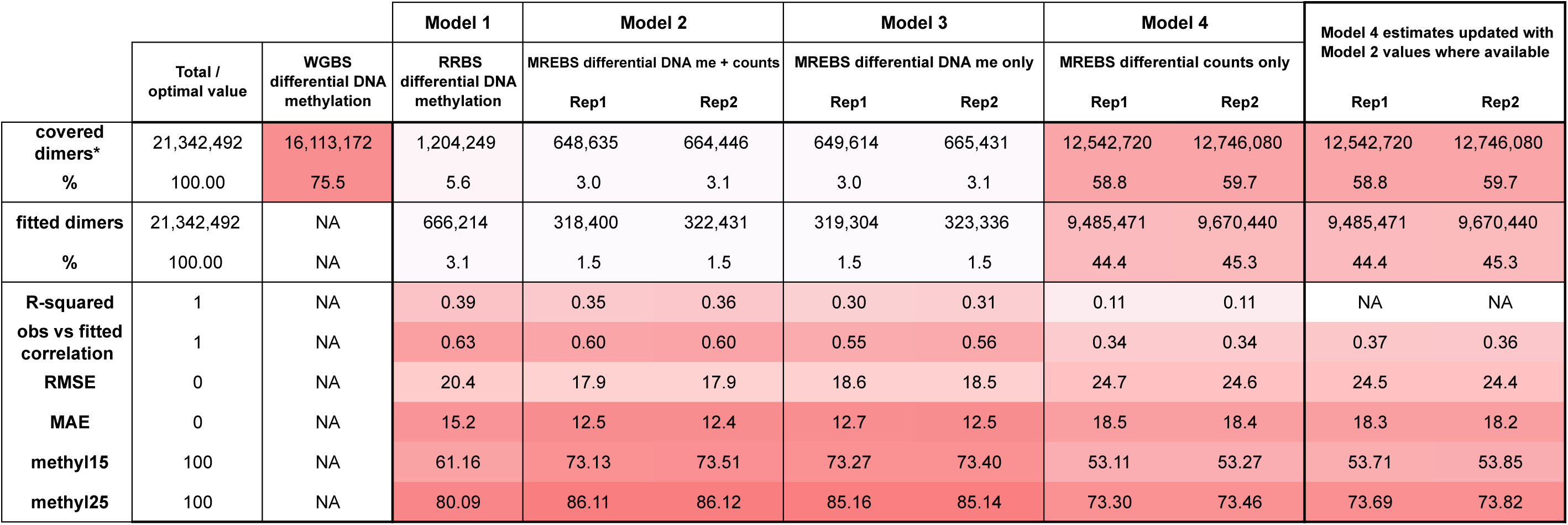
Differential DNA methylation model metrics. The table gives metrics (first column) for four different models, as well as a combined model, (top row and described in the text). The column labeled ‘Total / optimal value’ gives the maximum or best value achievable for each metric. The column labeled ‘WGBS differential DNA methylation’ provides coverage information for comparison purposes. *Note: In the case methylation levels, only CpG dimers with 5X coverage in both EARLY intermediates and ESCs were considered. With respect to counts, CpG dimes with 2+ reads in the surrounding 1Kb bin, in at least one sample, were considered. Red intensity signifies how close the metrics are to the optimal values.

Model 2 is a multiple regression model using both MREBS conversion-based differential DNA methylation (EARLY - ESC) and MREBS differential read count data to predict WGBS-based differential DNA methylation values. A model was built for each replicate pair. The fits for both replicates were similar (R2 = 0.35 and R2 = 0.36) and similar to those obtained using the RRBS data (Table 3). Interestingly, both RMSE (~18% for both replicates) and MAE (~12.5% for both replicates) values were better than those seen for the RRBS-based model. The concordance metrics (methyl15 = ~73% and methyl25 = ~86%, for both replicates) were also superior to that obtained for the RRBS-based model (Table 3). However, the differential DNA methylation values of only 1.5% (n = ~320 thousand) of CpG dimers genome-wide could be predicted, half of that predicted by the RRBS data (n = 666,214 or ~3% of CpG dimers). However, these percentages could be increased with greater sequencing depth.

Models 3 and 4 are both simple linear models using each of the two independent variables from model 2, respectively. The fit for model 3, using only MREBS conversion-based differential DNA methylation (EARLY - ESC), was somewhat worse without the additional count data (R2 = 0.30 and R2 = 0.31 for the replicate pairs), but, interestingly, the RMSE (~18.5% for both replicates) and MAE (~12.5% for both replicates) is still superior to that of the RRBS based model, as are the concordance metrics (methyl15 = ~73% and methyl25 = ~85%, for both replicates) (Table 3).

Model 4 is based only on MREBS differential read counts within 1kb window around each CpG (EARLY - ESC). Only those CpGs with at least two reads in the surrounding +/- 500bp in at least one sample were considered, amounting to 12.5 (58.8%) and 12.8 (59.7%) million CpG dimers for MREBS replicate 1 and 2, respectively (Table 3). This represents ~10X more CpG dimers than those that are available for use in the RRBS-based model 1 (1.2 million CpG dimers), and ~20X more CpG dimers than those that are available for use in models 2 and 3 based on MREBS sites with 5X coverage in both samples (~648–665 thousand CpG dimers). Although the fits for model 4 are worse than those based on the MREBS conversion-based DNA methylation estimates (R2 = 0.11 for both replicate pairs), the RMSE (~24–25%) and MAE (~18.5%) are not that much worse than the RRBS based model, nor are the concordance metrics (methyl15 = ~53% and methyl25 = ~73%) (Table 3). However, the differential DNA methylation values for 44–45% of CpGs were predicted using model 4, representing the overlap of those CpG dimers with 5X coverage by WGBS and those CpGs with at least 2 MREBS reads within the surrounding 1kb window.

To sum the benefit of both the extended coverage of model 4 and the improved accuracy of model 2, we combined their results, updating the model 4 estimates with those of model 2 where available. This marginally improved all the applicable metrics discussed previously (Table 3). Figure 4 shows how these combined differential DNA methylation predictions (iii, green tracks, two replicates) compared to WGBS (i, dark blue tracks) and RBBS (ii, light blue tracks) at different length scales: 611kb (A), 19kb (B), and an extended locus partitioned in three 18kb panels (C). Below the modeled estimates are tracks showing the MREBS conversion-based differential DNA methylation (iv, orange, two replicates) and MREBS-based differential read counts (v, red, two replicates) – the data that was combined. While the MREBS conversion-based differential DNA methylation coverage is comparable to that of the RRBS data (cf. tracks iv and ii, Table 3), the MREBS differential read count coverage approached that obtained using WGBS data (cf. tracks v and i, Table 3). In other words, the majority of the differential DNA estimates modeled on MREBS data are obtained by the read count data. These estimates track WGBS-based estimates, both for regions that are more methylated in the EARLY intermediates (Figures 4B/C and Supplementary Figure 1A), as well as regions that are more methylated in the ESCs (Figures 4A and Supplementary Figure 1B).

**Figure 4.**
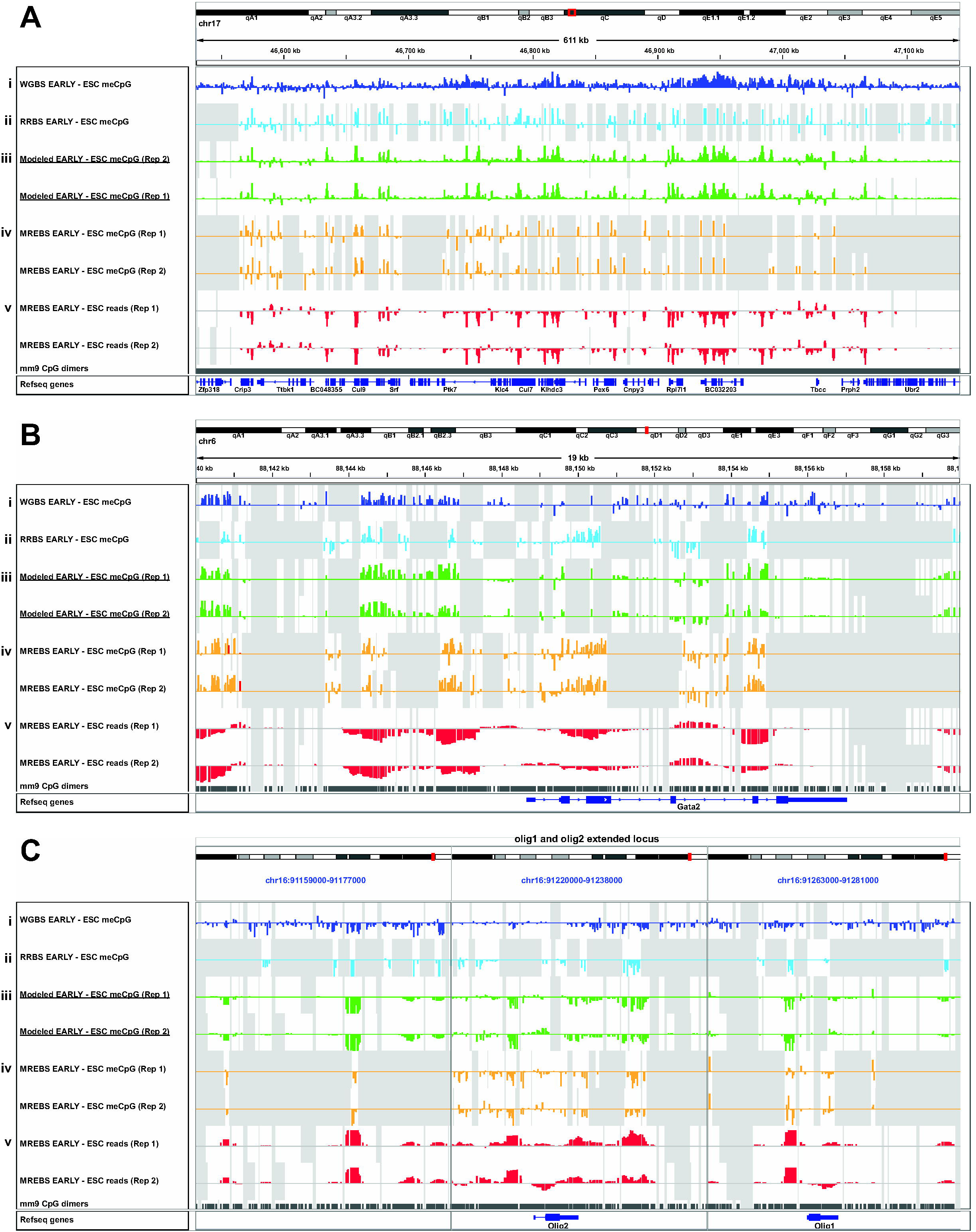
Differential DNA methylation levels modeled using MREBS data. A. IGV tracks of differential DNA methylation estimates between EARLY intermediates and ESCs (EARLY - ESC) based on WGBS data (i, dark blue), RRBS data (ii, light blue), modeled data based on combined model 2 (iii, green, two replicates), MREBS conversion-based DNA methylation estimates (iv, orange, two replicates), and MREBS read count-based estimates (v, red, two replicates), within a 611 kb region of chr17. Bottom two tracks show CpG dimer and Refseq gene locations. Gray background reflects regions (CpG dimers) where data was not available. B. As in (A), but for a 19kb region around the Gata2 gene. C. As in (A), but for the extend Olig1/2 gene locus, divided into three 18 kb panels.

## DISCUSSION

WGBS(22) and RRBS(23) are two popular bisulfite sequencing based methods for assessing DNA methylation levels. WGBS can potentially determine the methylation status of every single cytosine, but the amount of sequencing required to obtain sufficient coverage to do so can be beyond the scope of most projects. Sequencing demands are significantly reduced by using RRBS, but one incurs an 80–90% loss in the number of cytosines that can be measured. In order to address these respective shortcomings, we introduce methylation-sensitive restriction enzyme bisulfite sequencing (MREBS), which adds a bisulfite conversion step to the existing MRE protocol, methylation-sensitive restriction enzyme digestion followed by high-throughput sequencing (MRE-seq)(15).

Due to MREs reliance on methylation sensitive endonucleases, the distributions of the MREBS conversion-based DNA methylation estimates were systematically lower than those obtained using WGBS or RRBS data. However, the MREBS conversion-based estimates followed similar trends across all chromatin states (Figure 2). Moreover, high-confidence MREBS conversion-based DNA methylation estimates (CpGs with 5X coverage) were particularly enriched in chromatin states with the lowest DNA methylation levels (Figure 3). Since these chromatin states are known to be associated with gene regulation, their enrichment in MREBS data is beneficial. For MREBS libraries, 4.3–4.6% of CpG dimers had at least 5X read coverage, which we set as a threshold to generate bisulfite conversion based DNA methylation level calls. This fraction was comparable to the coverage obtained from RRBS libraries at a similar level of sequencing. However, for MREBS, ~60% of CpG dimers had two or more reads falling within the surrounding 1kb window in at least one of the two cell types (Supplementary Table 4). These can be used for estimating differential DNA methylation of a high proportion of CpGs, since MREBS utilizes methylation sensitive digestion and therefore read counts around CpGs anti-correlate with their methylation levels (Tables 5 and 6).

**Figure 3.**
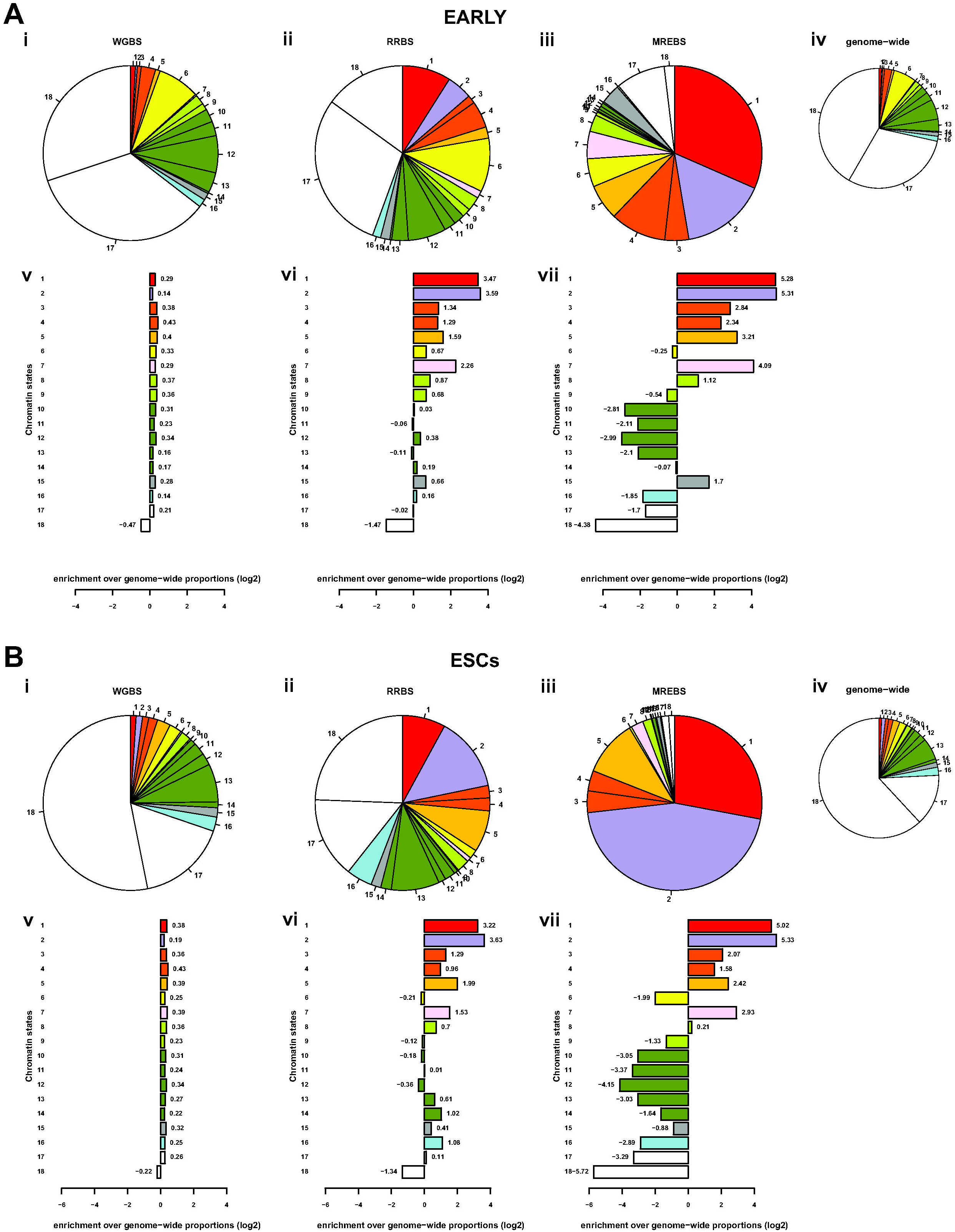
Chromatin state coverage by DNA methylation estimates by WGBS, RRBS, and MREBS. A. Pie charts show the proportion of 200 bp windows with DNA methylation estimates found in each of the 18 chromatin states (as described in Figure 2.2A) using i) whole-genome bisulfite sequencing (WGBS), reduced representation bisulfite sequencing (RRBS), and iii) methylation-sensitive restriction enzyme bisulfite sequencing (MREBS) EARLY intermediate samples, as compared to the proportion of chromatin states in genome for all 13.3 million windows (iv). Bar plots show the log2 fold change (observed / expected) number of windows with estimates per method: i) WGBS), RRBS, and iii) MREBS. The mean DNA methylation estimates within 200 bp windows corresponding to those used for the chromatin state model were used, only considering those windows containing at least one CpG with 5X coverage, in an effort to ensure high-confidence estimates. B. As in (A), but for ESCs.

To obtain estimates of differential DNA methylation based on MREBS data, we built a multiple regression model that incorporates both MREBS conversion fractions and read count data to predict differential DNA methylation values for ~3% of CpGs. The fits for both replicates were similar and correspondence metrics comparing the model-fitted values to WGBS estimates were superior to those obtained from models built using RRBS methylation data alone (Table 3). Differential DNA methylation estimates for a much greater proportion of CpGs (~60%) could be obtained using a model that used only MREBS differential read data within 1kb windows around CpG sites. The accuracy of MREBS read count-based models was lower than those based on conversion ratios, nonetheless, the dramatically higher coverage makes these data useful for low resolution differential methylation estimates (Table 3, Figures 4 and 5). In this study, we utilized 1kb windows around CpG dimers for this purpose, but one could use different windows, as one sees fit.

In summary, with respect to conversion-based DNA methylation estimates, MREBS provides a similar level of coverage to that obtained using RRBS. However, with MREBS one can additionally obtain DNA methylation estimates for a much larger proportion of the genome based on differential MREBS read counts around CpGs, providing a level of coverage that approaches that obtained by WGBS at a fraction of the cost.

## AUTHOR CONTRIBUTIONS

G.B. participated in project planning and data interpretation, performed bioinformatics analysis, and wrote the manuscript. M.M. and L.R. participated in project planning and data interpretation, and generated experimental data. C.C. produced experimental data. K.P. participated in data interpretation, provided supervision, and edited the manuscript. M.P. conceived the study, supervised the project, interpreted the data, provided guidance for bioinformatics analysis, and edited the manuscript.

## SUPPLEMENTARY DATA

Supplementary Data are available at NAR online.

## ACKNOWLEDGEMENTS

We thank Bernadett Papp for critical reading of this manuscript. We also thank the Broad Stem Cell Research Center High-Throughput Sequencing Core.

## TABLE AND FIGURES LEGENDS

**Supplementary Figure 1. Examples of modeled differential DNA methylation around gene loci**. A. As in Figure 4A, but for 13 genes up-regulated in EARLY intermediates relative to ESCs. B. As in Figure 4A, but for 10 genes up-regulated in ESCs relative to EARLY intermediates.

**Supplementary Table 1. Bisulfite sequencing library mapped reads and mean CpG coverage depth.** WGBS and MREBS reads were mapped the whole genome (mm9). Mean CpG coverage determined for CpGs on either strand. Mean CpG coverage determined for CpGs on either strand. RRBS reads were mapped to an in silico MspI digested reduced reference genome. MREBS reads were filtered in silico to have the expected 5’ start sites.

**Supplementary Table 2. MRE endonuclease recognition sequence frequency within the mm9 genome.** MRE endonuclease recognition sites are highlighted to show their position within the ranked frequencies for all the 4mer CpG, including chrM. *Note: HpaII has the same recognition sequence as MpsI (the endonuclease typically used for RRBS libraries), albeit HpaII is methylation sensitive, as are Acil and Hin6I.

**Supplementary Table 3. CpG dimer coverage per bisulfite sequencing library.** The percentage of CpG dimers with at least 5X coverage for each bisulfite sequencing library, as well as the percentage of CpG dimers with the specified number of MREBS reads within a surrounding 1kb window. *Note: This is based on 21,342,493 CpG dimers in mm9, excluding chrM.

**Supplementary Table 4. CpG dimer coverage for differential analysis per bisulfite sequencing library.** The percentage of CpG dimers with at least 5X coverage in both the EARLY intermediate and ESC samples for each bisulfite sequencing library, as well as the percentage of CpG dimers with at least two MREBS reads within a surrounding 1kb window. *Note: This is based on 21,342,493 CpG dimers in mm9, excluding chrM.

**Supplementary Table 5. Differential DNA methylation model coefficients.** Coefficient values (first column) for four different models (top row and described in the text).

